# RNA-protein interaction analysis of SARS-CoV-2 5’- and 3’-untranslated regions identifies an antiviral role of lysosome-associated membrane protein-2

**DOI:** 10.1101/2021.01.05.425516

**Authors:** Rohit Verma, Sandhini Saha, Shiv Kumar, Shailendra Mani, Tushar Kanti Maiti, Milan Surjit

## Abstract

Severe acute respiratory syndrome-coronavirus-2 (SARS-CoV-2) is a positive-strand RNA virus. Viral genome is capped at the 5’-end, followed by an untranslated region (UTR). There is poly-A tail at 3’-end, preceded by an UTR. Self-interaction between the RNA regulatory elements present within 5’- and 3’-UTRs as well as their interaction with host/virus-encoded proteins mediate the function of 5’- and 3’-UTRs. Using RNA-protein interaction detection (RaPID) assay coupled to liquid chromatography with tandem mass-spectrometry, we identified host interaction partners of SARS-CoV-2 5’- and 3’-UTRs and generated an RNA-protein interaction network. By combining these data with the previously known protein-protein interaction data proposed to be involved in virus replication, we generated the RNA-protein-protein interaction (RPPI) network, likely to be essential for controlling SARS-CoV-2 replication. Notably, bioinformatics analysis of the RPPI network revealed the enrichment of factors involved in translation initiation and RNA metabolism. Lysosome-associated membrane protein-2a (Lamp2a) was one of the host proteins that interact with the 5’-UTR. Further studies showed that Lamp2 level is upregulated in SARS-CoV-2 infected cells and overexpression of Lamp2a and Lamp2b variants reduced viral RNA level in infected cells and vice versa. In summary, our study provides an useful resource of SARS-CoV-2 5’- and 3’-UTR binding proteins and reveal the antiviral function of host Lamp2 protein.

**Importance:** Replication of a positive-strand RNA virus involves an RNA-protein complex consisting of viral genomic RNA, host RNA(s), virus-encoded proteins and host proteins. Dissecting out individual components of the replication complex will help decode the mechanism of viral replication. 5’- and 3’-UTRs in positive-strand RNA viruses play essential regulatory roles in virus replication. Here, we identified the host proteins that associate with the UTRs of SARS-CoV-2, combined those data with the previously known protein-protein interaction data (expected to be involved in virus replication) and generated the RNA-protein-protein interaction (RPPI) network. Analysis of the RPPI network revealed the enrichment of factors involved in translation initiation and RNA metabolism, which are important for virus replication. Analysis of one of the interaction partners of the 5’-UTR (Lamp2a) demonstrated its antiviral role in SARS-CoV-2 infected cells. Collectively, our study provides a resource of SARS-CoV-2 UTR-binding proteins and identifies an antiviral role of host Lamp2a protein.

## Introduction

In December 2019, a highly pathogenic coronavirus was identified in the Wuhan city, China, which was named as 2019-nCoV/SARS-CoV-2 (1,2,3). Since then, the virus has spread globally and World health organization (WHO) has classified the outbreak as a pandemic. SARS-CoV-2 belongs to the family of *Coronaviridae*. Coronaviruses are known to be present in animals and humans for a long time, usually resulting in respiratory and intestinal dysfunction in the host (4). Until the emergence of Severe acute respiratory syndrome coronavirus (SARS-CoV) in 2002, coronaviruses were not considered to be a significant threat to human health (5). However, within last 18 years since the SARS outbreak, three major human coronavirus outbreaks have occurred, suggesting that highly pathogenic human coronaviruses are evolving fast. Considering the severity of the current SARS-CoV-2 pandemic, there is an urgent need to understand the life cycle and pathogenetic mechanism of the virus. Since all coronavirus genomes share significant homology, the knowledge obtained from the study of SARS-CoV-2 genome will be useful to formulate a long-term action plan to deal with the current as well as future coronavirus outbreaks. Coronaviruses are positive strand RNA viruses. Viral genome serves as the template for synthesis of antisense strand, production of proteins involved in replication as well as assembly of the progeny virions (6). Viral genome is capped at the 5’-end and poly adenylated at the 3’-end. 5’ and 3’-ends of the genome contain noncoding sequences, also known as untranslated regions (UTRs). 5’- and 3’-UTRs in the positive strand of RNA viruses play essential regulatory roles in virus replication, enhancing the stability of the viral genomic RNA, host immune modulation and encapsidation of the viral genome into the nucleocapsid core. Additionally, cis-acting regulatory elements are present within the coding regions of the positive and negative strand (replication intermediate) of the viral genome. These regulatory RNA elements also play significant roles in the life cycle and pathogenesis of the virus. Intraviral interactions between regulatory RNA elements of the virus and intraviral as well as virus-host RNA-protein interactions control the function of the 5’-, 3’-UTRs and internal cis-acting RNA elements of the virus (7, 8).

SARS-CoV-2 contains a 260-nucleotide long 5’-UTR and a 200 nucleotide long 3’-UTR. These UTRs show considerable homology with the 5’- and 3’-UTRs of other beta coronaviruses such as SARS and SARS related beta coronaviruses (9). Distinct stem loops and secondary structures within the UTRs are known to mediate their regulatory function. Notably, stem loop I and stem loop II (SL-I and SL-II) of the 5’-UTR are important for long range interaction and subgenomic RNA synthesis, respectively. SL-III contains the translation-regulatory sequence (TRS), which is essential for the discontinuous transcription of ORF1ab. SL-5 is crucial for the viral RNA packaging and translation of the ORF1ab polyprotein (10, 11, 12).

The 3’-UTR of coronaviruses is important for viral replication (RNA synthesis and translation). It contains a hyper variable region (HVR) which, has been shown to be important for pathogenesis of mouse hepatitis virus (MHV). HVR contains a stem loop II-like motif (S2M) that associates with host translation factors (13). S2M motif is also present in the 3’-UTR of SARS-CoV-2 (9).

Interaction of host proteins with the 5’- and 3’-UTRs of viral genomic RNA is important for replication and pathogenesis of many RNA viruses. Interaction of polypyrimidine tract binding protein (PTB) with the 5’-UTR-TRS of MHV is known to control transcription of the viral RNA (14, 15, 16). Further, RNA helicases such as DDX1, DHX15; proteins involved in translation regulation such as eIF1*α*, eIF3S10; proteins involved in cytoskeleton movement such as tubulin, Annexin A2, moesin as well as GAPDH (glyceraldehyde 3-phosphate dehydrogenase) is known to associate with the 5’-UTR of few coronaviruses (7, 17). 3’-UTR of MHV associates with hnRNPA1 and modulate viral replication (18). Poly-A binding protein (PABP), transcriptional activator p100 (SND1), heat shock proteins (HSP40, HSP 60, HSP 70) associate with the 3’-UTR of MHV (19). Functional significance of some these interactions are known (8).

Despite general acceptance of the importance of the 5’- and 3’-UTRs of RNA viruses in controlling their replication and pathogenesis, no systematic study has been under taken to elucidate the molecular composition of the RNA-protein complex assembled at the 5’- and 3’-UTRs of SARS-CoV-2 and function of 5’- and 3’-UTRs of SARS CoV-2. We employed RNA-protein interaction detection (RaPID) assay coupled to liquid chromatography with tandem mass spectrometry (LC-MS-MS) to identify the repertoire of host proteins that interact with the SARS-CoV-2 5’- and 3’-UTR (20). These datasets were used to construct the virus-host RNA-Protein-protein interaction (RPPI) network. *In silico* analyses of the RPPI network revealed enrichment of proteins involved in multiple processes such as Cap-dependent translation and RNA metabolism. Further studies revealed an antiviral role of LAMP2 during SARS-CoV-2 infection. Functional significance of these findings in SARS-CoV-2 replication and pathogenesis is discussed.

## Results

### Identification of host proteins that interact with the 5’- and 3’-UTRs of SARS-CoV-2 genomic RNA

300 nucleotides from the 5’-end (denoted as 5’-300 RNA) and 203 nucleotides form the 3’-end (3’-203 RNA) of the SARS-CoV-2 genomic RNA (Wuhan isolate), which includes the 5’- and 3’-UTRs, respectively (nucleotides 1-264 and 29676-29878 in the viral genome), were cloned into the pRMB vector in between the BirA Ligase binding stem loop (SL-A and SL-B) sequences (Fig 1A-C). 5’- and 3’-UTRs consist of 264 and 228 nucleotides, respectively. Extra bases were included at the 5’-end to ensure that stem loops in the predicted secondary structures of the 5’-UTR remain intact in the hybrid RNA (Fig 1A). Eight Adenine residues were retained at the 3’-end (Fig 1B). “mfold” mediated comparison of the secondary structures of the 5’-300 and 3’-203 RNA sequences with and without BirA ligase binding stem loop RNA sequences indicated that secondary structures of viral 5’-UTR and 3’-UTR and BirA binding stem loops (SL-A and SL-B) are not disturbed in the hybrid RNA sequence (compare SL-I to SL-Vc in the 5’-300 RNA and SL-A and SL-B in Fig 1A and S1 Fig; compare SL-I to SL-IV and SL-A and SL-B in Figure 1B and S2 Fig). RaPID (RNA-Protein interaction detection) assay was performed to identify the host proteins that interact with the 5’-300 and 3’-203 RNA in HEK293T cells. RaPID assay allows BirA ligase mediated *in vivo* biotin labeling of host proteins that interact with the RNA sequence of interest cloned in between the BirA ligase-binding RNA motifs. Biotinylated proteins are captured using streptavidin agarose beads and identified by LC-MS-MS (Fig 1C, 20). Thus, both transiently interacting and stably interacting proteins can be identified using this technique.

**Figure 1.**
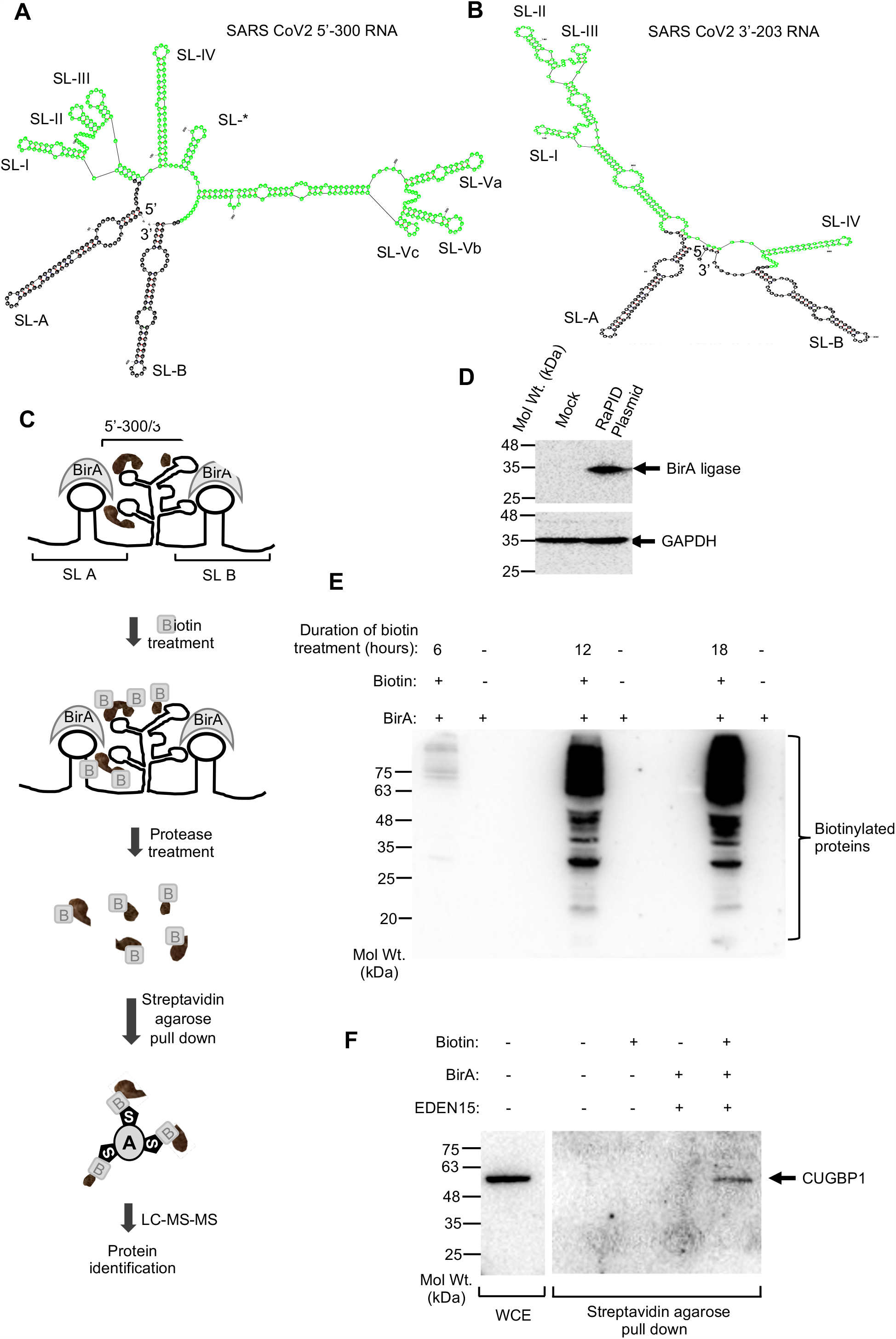
Establishment of RaPID assay to identify the host interaction partners of SARS-CoV-2 5’-300 and 3’-203 RNA. A. Schematic of predicted secondary structure of SARS-CoV-2 5’-300 RNA. “*” denotes an unannotated stem loop in the 5’-UTR. B. Schematic of predicted secondary structure of SARS-CoV-2 3’-203 RNA. C. Schematic of RaPID assay workflow. Biotin is represented as ‘B’, agarose is represented as ‘A’ and streptavidin is represented as ‘S’. D. Western blot detection of BirA ligase and GAPDH level in HEK293T cells transfected with the RaPID plasmid for 48 hours. E. Western blot detection of Biotinylated protein in HEK293T cells transfected with BirA ligase and treated with Biotin for different time points, as indicated. F. Western blot detection of CUGBP1 protein in the whole cell extract of HEK293T cells (WCE) and in streptavidin agarose pull down of EDEN 15 RNA interacting biotinylated proteins.

Expression of BirA ligase in HEK293T cells was confirmed by western blot using anti-HA antibody that recognizes the HA-tag fused to the BirA ligase (Fig 1D). Optimal duration of biotin treatment was selected through a time course analysis of biotin labeling, based on which 18 hour labeling period was selected (Fig 1E). To test the functionality of the RaPID assay, interaction between CUGBP1 protein and EDEN15 RNA was tested in the HEK293T cells (20). Streptavidin pulldown of EDEN15 RNA interacting proteins, followed by western blot using CUGBP1 antibody revealed the interaction between them, in agreement with earlier report (Fig 1F, 20).

HEK293T cells expressing the SL-A-5’-300-SL-B or SL-A-3’-203-SL-B or SL-A-SL-B (pRMB) was treated with Biotin, followed by pulldown of biotinylated proteins and LC-MS-MS analysis. Biotin untreated cells were processed in parallel as control. Samples from three independent experiments were run in triplicate. Pearson correlation analysis of MS data demonstrated good correlation between replicates for individual biological samples (average range: 0.5-0.98, Fig 2A). Specific and strong interaction partners of 5’-300 and 3’-203 RNAs were identified in three steps (a) only those proteins having at least one biotinylated peptide and a “Pep score” of 15 or more in all LC-MS samples were selected [(S3 Fig, S1 Table) (21]; (b) subtraction of pRMB dataset from 5’-300 and 3’-203 datasets (Fig 2B); (c) from the background subtracted dataset, only those proteins with minimum 2 unique peptides and “Prot score” of 40 or more were considered for further analysis (Table 1).

**Table 1:**
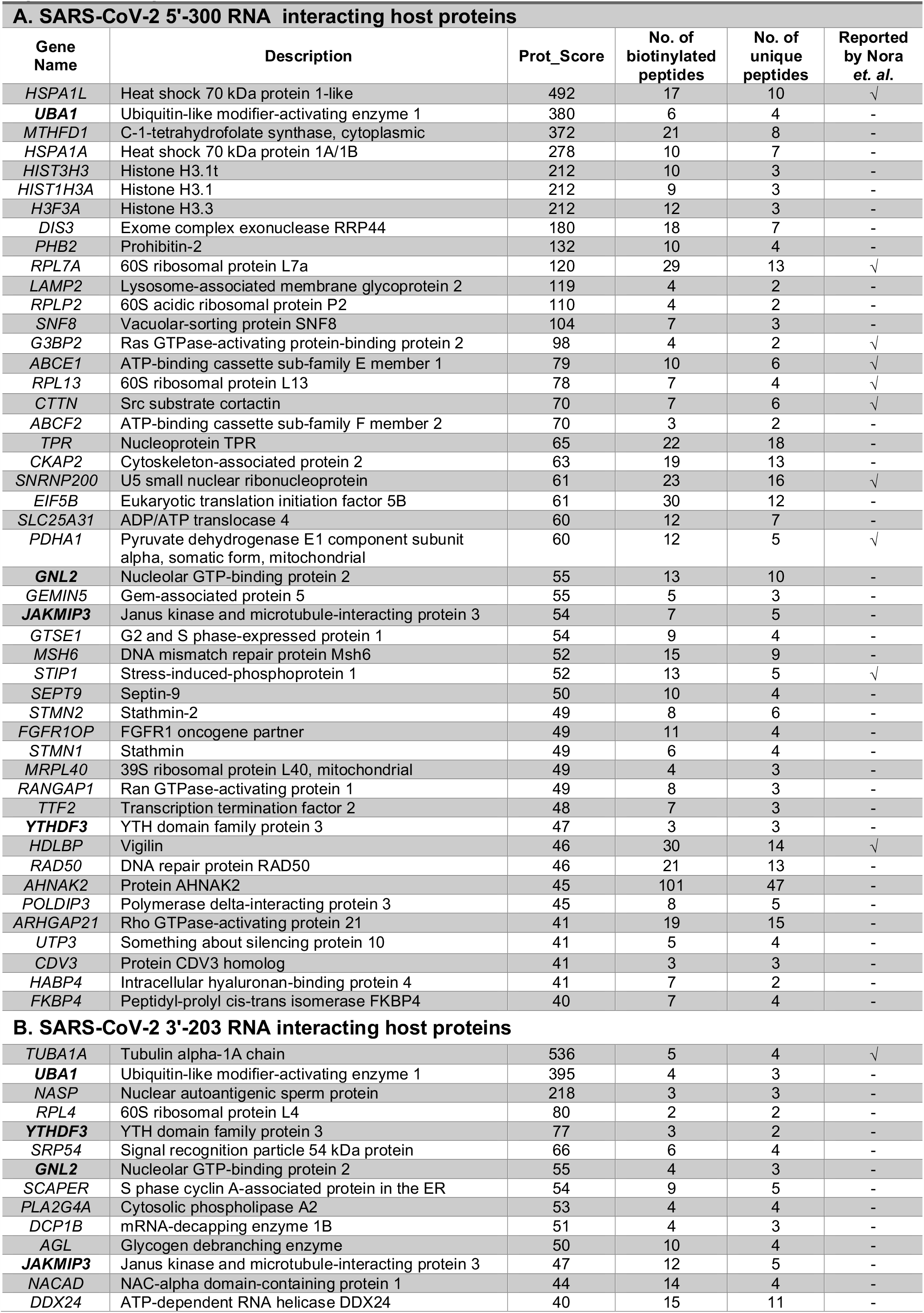
Host proteins that interact with the 5’-300 and 3’-203 RNA of SARS-CoV-2, identified by RaPID assay.

**Figure 2.**
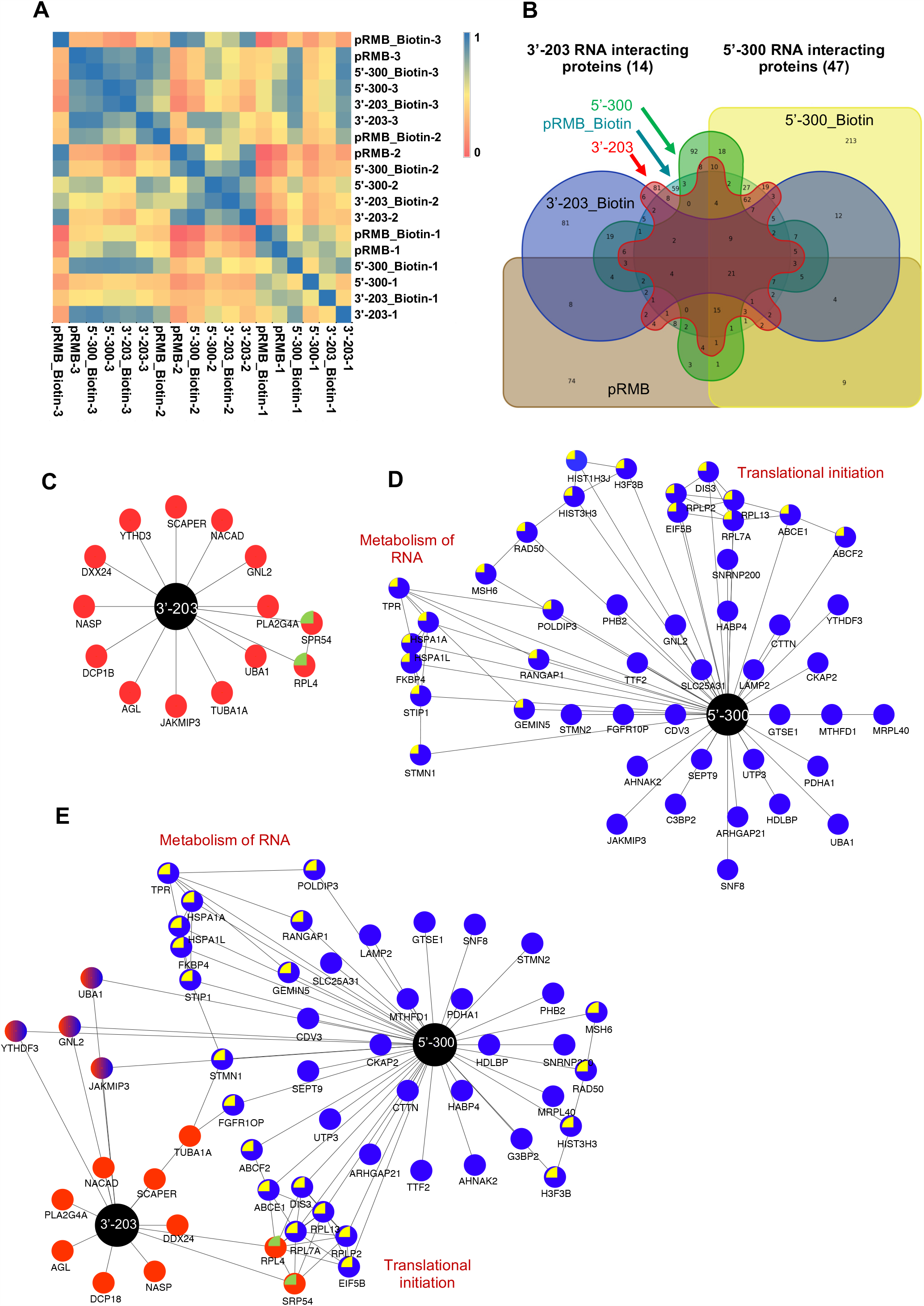
Identification of SARS-CoV-2 5’-300 and 3’-203 RNA interacting host proteins by RaPID assay. A. Clustering analysis of correlation between biological replicates used in LC-MS-MS. B. Venny analysis of 5’-300 and 3’-203 RNA interacting proteins. (3’-203 in red, 3’-203_Biotin in blue, 5’-300 in green, 5’-300_Botin in yellow, pRMB in brown and pRMB_Biotin in dark green). C. Schematic of RNA-protein interaction network of the 3’-203 SARS-CoV-2 RNA. Black node: 3’-203; Red node: Host protein. Host proteins that interact with each other contain green color inside the node. D. Schematic of RNA-protein interaction network of the 5’-300 SARS-CoV-2 RNA. Black node: 5’-300; Blue node: Host protein. Host proteins that interact with each other contain yellow color inside the node. E. Schematic of RNA-protein interaction network of the 5’-300+3’-203 SARS-CoV-2 RNAs. Black nodes: 5’-300 or 3’-203; Blue or red nodes: Host protein. Host proteins that interact with each other contain yellow or green color inside the node. Common host proteins that interact with both 5’-3000 and 3’-203 RNA are represented with dual (red+blue) color.

Following similar protocol, in an unrelated study, we had identified host interaction partners of the 3’-end (3’-UTR + adjacent 100 nucleotides) of the Hepatitis E virus (HEV) genomic RNA (S2 Table). HEV is a positive strand RNA virus of the *hepeviridae* family and 3’-end of the HEV harbors a secondary structure composed of multiple stem loops that have been predicted to be important for viral replication (22). In order to further increase our confidence in reliability of the RaPID technique, HEV 3’-end RNA interacting protein list was compared to the SARS-CoV-2 5’-300 and 3’-203 RNA-interacting protein list. One protein (Histone H3.1) was found to be common between the two datasets, indicating specificity of the interacting proteins for the RNA baits. Therefore, using RaPID assay, 47 and 14 host proteins were identified to interact with the 5’-300 and 3’-203 RNA, respectively (Table 1). Out of these, 4 proteins (UBA1, GNL2, JAKMIP3 and YTHDF3) interacted with both 5’- and 3’-UTRs (Table 1, represented in bold font). A recent publication reported the atlas of proteins bound to the SARS-CoV-2 RNA using the RAP-MS (RNA Antisense Purification with Mass Spectrometry) technique (23). Comparison with that data identified 10 proteins to be present in our 5’-300 dataset and 1 protein in the 3’-203 RNA binding protein data set (Table 1). Based on human protein atlas data, except HIST3H3 and SLC25A31, rest 55 proteins are expressed in both human lungs and intestine (S3 Table).

### Construction and analysis of the RNA-Protein-Protein interaction network at the 5’- and 3’-ends of the SARS-CoV-2 genome

5’-300 and 3’-203 RNA binding protein datasets were imported to “Cytoscape” to construct the RNA-Protein-protein interaction networks (RPPI network) (24). 3’-203 interacting proteins did not show any significant network characteristics (observed number of edges =1, expected no. of edges =1, numbers of node =14 Average node degree = 0.143, average clustering coefficient = 0.143 and PPI enrichment P value = 0.437) (Fig 2C). However, 5’-300 and 5’-300+3’-203 interaction RPPI networks showed significant enrichment of interactions [(5’-300 RPPI network characteristics: observed number of edges =27, expected no. of edges =10, numbers of node =46, average node degree = 1.17, average clustering coefficient = 0.311 and PPI enrichment P value = 3.64e-06); (5’-300+3’-203 RPPI network characteristics: observed number of edges =39, expected no. of edges =14, numbers of node =56, average node degree = 1.39, average clustering coefficient = 0.29 and PPI enrichment P value = 1.27e-08)]. Note that, based on the above network parameters, 5’-300+3’203 RPPI network is stronger than the 5’-300 network (Fig 2D, 2E).

Recently protein interaction map of SARS-CoV-2 has been reported (25). Since RNA-protein complex assembled at the 5’- and 3’-UTR as well as Viral RNA-dependent RNA polymerase (RdRp)-associated protein complex are known to drive the viral replication process in positive strand RNA viruses, we combined the 5’-300+3’-203 RPPI dataset and previously reported PPI dataset (only the subset of PPI proposed by Gordon and coworkers to be involved in viral replication) to generate an integrated RPPI network of SARS-CoV-2 replication machinery (S4 Fig). As expected, the integrated RPPI network showed significantly higher PPI enrichment compared to replication PPI alone [(Integrated RPPI network characteristics: observed no. of edges =328, expected no. of edges =152, number of nodes =196, average node degree = 3.35, average clustering coefficient = 0.492 and PPI enrichment P value = 1.0e-16); (replication PPI network characteristics: observed no. of edges =191, expected no. of edges =78, numbers of nodes = 143, average node degree = 2.67, average clustering coefficient = 0.501 and PPI enrichment P value = <1.0e-16)] (S4A and S4B Figs).

Next, gene ontology (GO) and Reactome pathway analysis of the RPPI network was performed using “GSEA tool” to determine the significantly enriched processes/pathways (S5 and S6 Figs) (26, 27). Of note, proteins involved in intracellular transport, translation initiation, amide biosynthetic pathway and mRNA metabolism were enriched in the GO-Biological processes category (Table 2A). Similarly, proteins involved in cellular response to external stimuli, HSP 90 chaperones for steroid hormone receptors, RNA metabolism, translation, Influenza infection, infectious disease, cell cycle etc. were enriched in Reactome pathway analysis (Table 2A).

**Table 1:** Gene ontology and Reactome pathway analyses of SARS-CoV-2 5’-300 and 3’-203 RNA-protein interaction network and identification of potential drug targets.

| Gene Set Name | Observed gene count | p-value | Gene names |
| --- | --- | --- | --- |
| **Gene Ontology (Biological Process)** |  |  |  |
| GO_INTRACELLULAR_TRANSPORT | 19 | 1.87E-12 | TPR, ABCE1, RPL7A, RPLP2, RPL13, RPL4, POLDIP3, SRP54, LAMP2, CTTN, PHB2, SNF8, HSPA1L, GTSE1, RANGAP1, NACAD, GEMIN5, TUBA1A, ARHGAP21 |
| GO_INTRACELLULAR_PROTEIN_TRANSPORT | 16 | 5.32E-12 | TPR, ABCE1, RPL7A, RPLP2, RPL13, RPL4, POLDIP3, SRP54, LAMP2, CTTN, PHB2, SNF8, HSPA1L, GTSE1, RANGAP1, NACAD |
| GO_TRANSLATIONAL_INITIATION | 9 | 8.13E-12 | TPR, ABCE1, RPL7A, RPLP2, RPL13, RPL4, YTHDF3, HABP4, EIF5B |
| GO_AMIDE_BIOSYNTHETIC_PROCESS | 14 | 2.51E-11 | TPR, ABCE1, RPL7A, RPLP2, RPL13, RPL4, POLDIP3, GEMIN5, YTHDF3, HABP4, EIF5B, MRPL40, PDHA1, MTHFD1 |
| GO_PROTEIN_CONTAINING_COMPLEX_SUBUNIT_ORGANIZATION | 18 | 1.38E-10 | TPR, ABCE1, SRP54, LAMP2, CTTN, GEMIN5, MRPL40, HSPA1A, SNRNP200, G3BP2, STMN2, STMN1, FKBP4, H3-3A, CKAP2, NASP, H3C1, H3-4 |
| GO_MRNA_METABOLIC_PROCESS | 13 | 2.80E-10 | RPL7A, RPLP2, RPL13, RPL4, POLDIP3, GEMIN5, YTHDF3, HABP4, HSPA1A, SNRNP200, DCP1B, DIS3, TTF2 |
| GO_PEPTIDE_BIOSYNTHETIC_PROCESS | 12 | 6.54E-10 | TPR, ABCE1, RPL7A, RPLP2, RPL13, RPL4, POLDIP3, GEMIN5, YTHDF3, HABP4, EIF5B, MRPL40 |
| GO_CELLULAR_AMIDE_METABOLIC_PROCESS | 14 | 9.20E-10 | TPR, ABCE1, RPL7A, RPLP2, RPL13, RPL4, POLDIP3, GEMIN5, YTHDF3, HABP4, EIF5B, MRPL40, PDHA1, MTHFD1 |
| GO_CELLULAR_MACROMOLECULE_LOCALIZATION | 17 | 1.32E-09 | TPR, ABCE1, RPL7A, RPLP2, RPL13, RPL4, POLDIP3, SRP54, LAMP2, CTTN, PHB2, SNF8, HSPA1L, GTSE1, RANGAP1, NACAD, SEPTIN9 |
| GO_PEPTIDE_METABOLIC_PROCESS | 12 | 4.67E-09 | TPR, ABCE1, RPL7A, RPLP2, RPL13, RPL4, POLDIP3, GEMIN5, YTHDF3, HABP4, EIF5B, MRPL40 |
| **Reactome pathway** |  |  |  |
| REACTOME_CELLULAR_RESPONSES_TO_EXTERNAL_STIMULI | 14 | 1.40E-13 | HSPA1A, RPL4, RPL7A, RPL13, RPLP2, TPR, TUBA1A, FKBP4, HSPA1L, STIP1, H3C1, H3-3A, H3-4, RAD50 |
| REACTOME_METABOLISM_OF_RNA | 12 | 1.60E-10 | HSPA1A, RPL4, RPL7A, RPL13, RPLP2, TPR, DIS3, UTP3, GEMIN5, DCP1B, POLDIP3, SNRNP200 |
| REACTOME_HSP90_CHAPERONE_CYCLE_FOR_STEROID_HORMONE_RECEPTORS_SHR | 5 | 1.93E-08 | HSPA1A, TUBA1A, FKBP4, HSPA1L, STIP1 |
| REACTOME_INFLUENZA_INFECTION | 6 | 1.01E-07 | HSPA1A, RPL4, RPL7A, RPL13, RPLP2, TPR |
| REACTOME_TRANSLATION | 7 | 2.11E-07 | RPL4, RPL7A, RPL13, RPLP2, SRP54, EIF5B, MRPL40 |
| REACTOME_INFECTIOUS_DISEASE | 10 | 3.64E-07 | HSPA1A, RPL4, RPL7A, RPL13, RPLP2, TPR, TUBA1A, H3C1, RANGAP1, SNF8 |
| REACTOME_RRNA_PROCESSING | 6 | 4.86E-07 | RPL4, RPL7A, RPL13, RPLP2, DIS3, UTP3 |
| REACTOME_CELL_CYCLE | 9 | 5.64E-07 | TPR, TUBA1A, H3C1, H3-3A, H3-4, RAD50, RANGAP1, CEP43, GTSE1 |
| REACTOME_SRP_DEPENDENT_COTRANSLATIONAL_PROTEIN_TARGETING_TO_MEMBRANE | 5 | 6.08E-07 | RPL4, RPL7A, RPL13, RPLP2, SRP54 |
| REACTOME_EUKARYOTIC_TRANSLATION_INITIATION | 5 | 8.19E-07 | RPL4, RPL7A, RPL13, RPLP2, EIF5B |

**B. Known drugs targeting SARS-CoV-2 5'-300 and 3'-203 RNA interacting host proteins**
| Viral RNA binding protein | Gene name | Drug name |
| --- | --- | --- |
| 5'-300 RNA interacting host proteins | RPL13 | Thalidomide |
|  | PDHA1 | NADH |
|  | HABP4 | Hyaluronan |
|  | MTHFD1 | Tetra hydro folic acid |
| 3'-203 RNA interacting host proteins | TUBA1A | Albendazole, Vinblastine, Mebendazole |
|  | PLA2G4A | Fluticasone Propionate |

Since, 5’- and 3’-UTRs are well conserved, drugs directly targeting the UTRs or acting upon the proteins that interact with the UTRs may function as potent antivirals by inhibiting the viral replication process. A search for known drugs against the 57 RNA binding proteins listed in Table 1 identified six proteins (RPL13, PDHA1, HABP4, MTHFD1, TUBA1A, PLA2G4A), against whom known drugs exist (Table 2B). Notably, Thalidomide, which targets RPL13, is being tested in clinical trial to evaluate its efficacy as an anti-inflammatory therapeutic in COVID-19 patients (28). C-1-tetrahydrofolate synthase (MTHFD1) is the target of Tetrahydrofolic acid or vitamin B9, which is being used as a supplement in CoVID-19 treatment (29). Phospholipase A2 (PLA2G4A) is the target of steroid drug Fluticasone propionate. A recent report shows the ability of another steroid drug “Ciclesonide” to block SARS-CoV-2 replication by targeting its replication-transcription complex (30).

### Lamp2 is a host restriction factor against SARS CoV-2

Lamp2a (CD107b) was identified as an interaction partner of the 5’-300 RNA in RaPID assay. LAMp2a is the receptor for chaperone mediated autophagy and recent studies have proposed a role of autophagy in SARS-CoV-2 pathogenesis (31). Hence, significance of the interaction between 5’-300 RNA and Lamp2a in SARS-CoV-2 life cycle was investigated using mammalian cell culture-based infection model of SARS CoV-2.

Vero E6 cell based infection model of SARS-CoV-2 was established to investigate the role of Lamp2a in SARS-CoV-2 life cycle. Productive infection of Vero E6 cells with SARS-CoV-2 was detected by immunofluorescence staining of the viral nucleocapsid (N) protein at 48 hour post-infection using anti-N antibody (Fig 3A). N-staining was not observed in uninfected cells (Fig 3A). Expression of N protein was also detected by western blot using anti-N antibody, 48-hour post infection (Fig 3B, upper panel). An aliquot of the samples were immunoblotted with anti-GAPDH antibody to monitor equal loading of protein in uninfected and infected cells (Fig 3B, lower panel). Next, level of viral RNA in the culture media (from released virus) and inside the cells (intracellular) was measured by the quantitative real time PCR (QRT-PCR) analysis of 48 hour and 72 hour SARS-CoV-2 infected samples using SYBR green based method. QRT-PCR detection of RNA polymerase II (RP II) and RNase P (RP) served as normalization control for intracellular and secreted RNA quantity, respectively. An increase in viral RNA level was observed in 72 hour infected intracellular and secreted samples, compared to 48 hour infected sample, indicating productive infection of Vero E6 cells with SARS-CoV-2 (Fig 3C). Aliquots of the RNA from culture media was also used in Taqman probe based one step QRT-PCR analysis using primers and probes recommended by the Centre for disease control (CDC), USA, for detection of SARS-CoV-2 (Fig 3D). Both SYBR green and Taqman QRT-PCR methods showed comparable result, hence SYBR green method was followed in subsequent experiments. Collectively, these data demonstrate the establishment of Vero E6 infection model of SARS-CoV-2 in our experimental setups.

**Figure 3.**
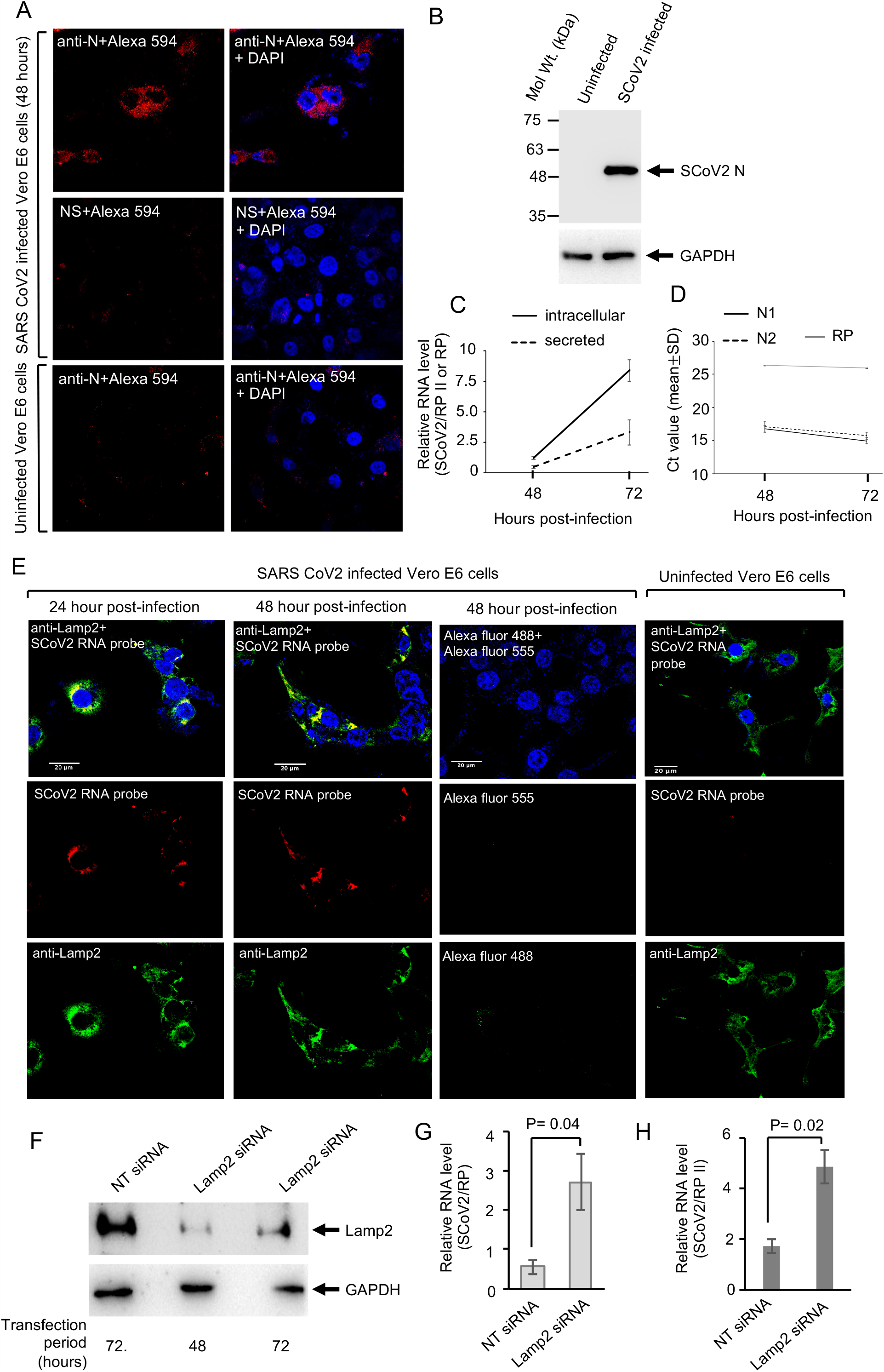
Lamp2 associates with the 5’-end of the SARS-CoV-2 genome and modulates the level of viral RNA in infected Vero E6 cells. A. Immunofluorescence staining of Nucleocapsid Protein (red) and nucleus (blue) in SARS-CoV-2 infected Vero E6 cells, 48 hour post-infection. B. Western blot detection of Nucleocapsid protein (upper image) and GAPDH (lower image) in SARS-CoV-2 infected Vero E6 cells, 48 hour post-infection. C. Level of SARS-CoV-2 RNA [normalized to that of RNase P (RP)] in the culture medium (secreted) and inside Vero E6 cells (intracellular) that are infected with the SARS-CoV-2 for the indicated periods. Real-time PCR reactions were performed using SYBR green based protocol. Data are mean ± SEM. D. Intracellular level of SARS-CoV-2 RNA in Vero E6 cells that are infected with SARS-CoV-2 for the indicated periods. Real-time PCR reactions were performed using Taqman probe-based protocol. N1, N2 and RP represents two regions within the SARS-CoV-2 Nucleocapsid coding region and one region within RNase P (RP) that was selected for PCR amplification. Data are mean ± SEM. E. Fluorescence *in situ* hybridization of SARS-CoV-2 5’-300 RNA and Lamp2, at indicated periods, post-infection. Lamp2 protein, SARS-CoV-2 RNA and Nucleus is denoted by green, red and blue, respectively. Scale bar is 20μM. F. Western blot detection of Lamp2 (Lamp2 antibody, upper panel) and GAPDH (lower panel) proteins in Vero E6 cells transfected for 72 hours with non targeting (NT) SiRNA or Lamp2 siRNA. G. Level of SARS-CoV-2 RNA (normalized to that of RP) in the culture medium of Vero E6 cells transfected with NT siRNA or Lamp2 siRNA and infected with SARS-CoV-2 for 48 hours. Data are mean ± SEM. H. Intracellular level of SRS CoV2 RNA (normalized to that of RP II) in Vero E6 cells treated with Lamp2 SiRNA and infected with SARS-CoV-2 for 48 hours. Data are mean ± SEM.

A fluorescence *in situ* hybridization (FISH) was conducted to monitor the interaction between 5’-300 region of viral genome and Lamp2 in SARS-CoV-2 infected Vero E6 cells. Lamp2 (green) and 5’-300 RNA specific probe (red) colocalized (indicated by yellow), suggesting their interaction (Fig 3E). Next, smart pool SiRNA targeting Lamp2 RNA was used to deplete Lamp2 protein in veroE6 cells. Western blot of Vero E6 whole cell extract after 48 and 72 hours of Lamp2 siRNA transfection demonstrated that siRNA was effective in reducing total Lamp2 level by >90% in siRNA transfected cells (Fig 3F, upper panel). GAPDH was used as a loading control (Fig 3F, lower panel). QRT-PCR of 72 hour siRNA transfected SARS-CoV-2 infected Vero E6 cells (analyzed 48 hour post-infection) revealed an increase in viral RNA level in both culture medium (Fig 3G) and inside the cells (Fig 3H).

Results obtained in Vero E6 cells were further verified in a human hepatoma (huh7) cells-based infection model of SARS CoV-2. Recent reports have shown the utility of Huh7 cells as a viable model for SARS-CoV-2 infection (23, 32). Infection of Huh7 cells by SARS-CoV-2 was confirmed by measuring the level of viral RNA inside the cells (intracellular) as well as in the culture medium (secreted). As expected, an increase in viral RNA level at 72 hour post-infection period (compared to 48 hour post-infection) indicated productive infection of these cells (S7A Fig). Further, western blot using anti-N antibody confirmed N-expression in infected samples (72 hour, S7B Fig). Lamp2 siRNA transfection reduced the corresponding protein level by >90% in Huh7 cells (S7C Fig). A significant increase in viral RNA level was observed in both culture medium and intracellular samples in 72 hour Lamp2 siRNA transfected and 48 hour SARS-CoV-2 infected Huh7 cells (S7D and 7E Figs).

Lamp2 is present in three different forms in mammalian cells that include Lamp2a, Lamp2b and Lamp2c, produced through alternative splicing (33). Lamp2a and Lamp2b are the major forms. Lamp2a is highly expressed in lungs, liver and placenta. Lamp2b varies from Lamp2a in the last 11 amino acids at its C-terminal sequence and it is highly expressed in Skeletal muscle (34). Lamp2 antibody used in this study recognizes all three Lamp2 variants. Next, the effect of overexpression of each Lamp2 isoform on SARS-CoV-2 replication was tested in Vero E6 cells. Overexpression of Lamp2a and Lamp2b significantly reduced the viral RNA level whereas Lamp2c overexpression did not alter viral RNA level at 48 hour post-infection, both in culture medium (Fig 4A) and intracellular samples (Fig 4B). Lamp2a, b and c overexpression was confirmed in aliquots of the sample using anti-lamp2 antibody, which recognizes all Lamp2 variants (Figs 4C-4E) as well as using anti-HA antibody (for Lamp2a). Similar results were obtained in Huh7 cell-based model of SARS-CoV-2 infection (S7F-7J Figs). Note that overexpressed Lamp2a and Lamp2b could be detected using both anti-HA and anti-Lamp2 antibodies in Huh7 cells but overexpressed Lamp2b could be detected using only anti-Lamp2 antibody in Vero E6 cells. Also note that no epitope tag is present in Lamp2c expression clone. Hence it was detected using anti-Lamp2 antibody.

**Figure 4.**
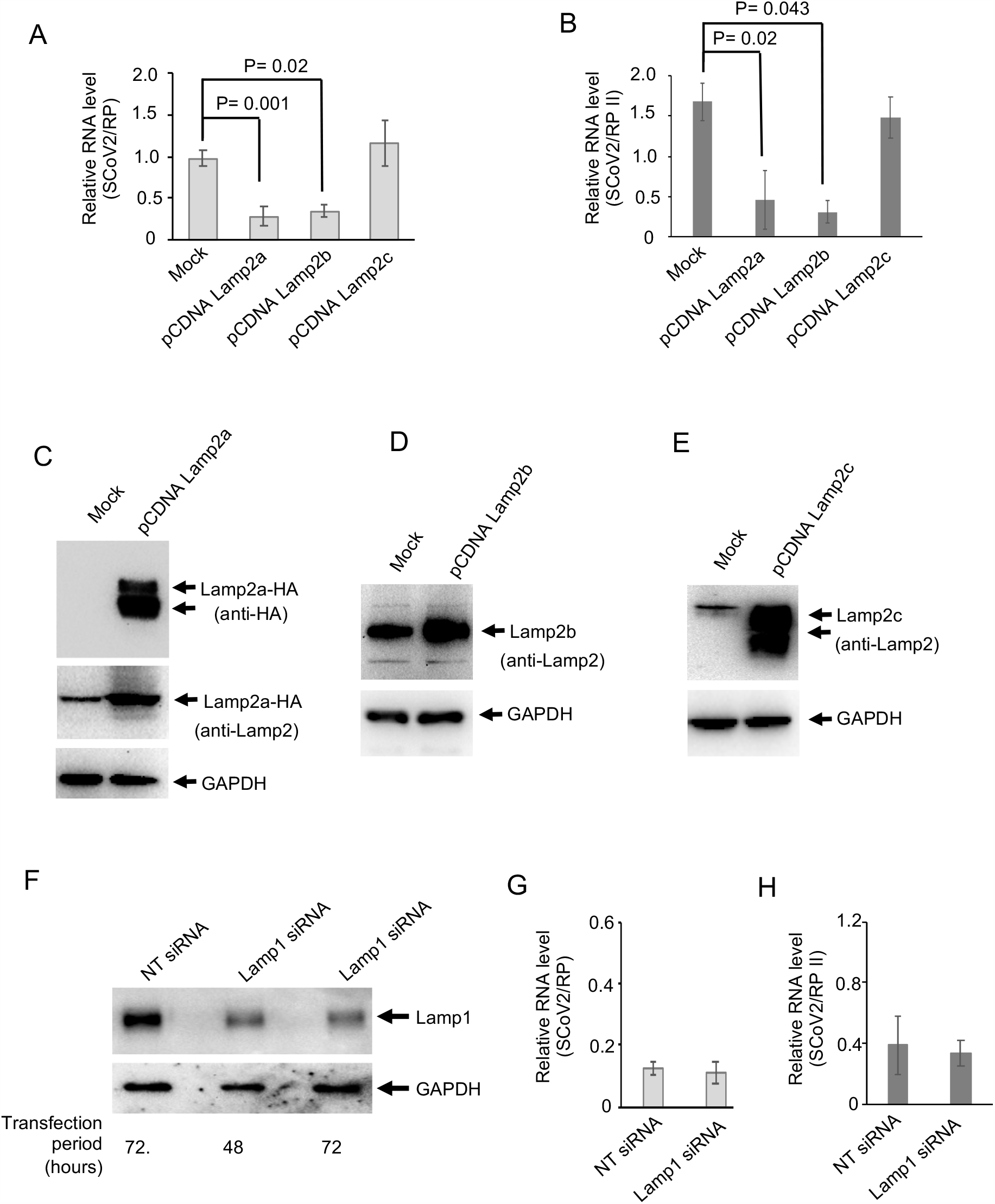
Lamp2 modulates viral RNA level in SARS-CoV-2 infected cells. A. Western blot detection of Lamp2a protein level [using anti-HA antibody (upper panel) and anti-Lamp2 antibody (middle panel)] and GAPDH protein level (lower panel) in mock transfected or pcDNA Lamp2a transfected Vero E6 Cells. B. Western blot detection of Lamp2b protein level using anti-Lamp2 antibody (upper panel) and GAPDH protein level (lower panel) in mock transfected or pcDNA Lamp2b transfected Vero E6 Cells. C. Western blot detection of Lamp2c protein level using anti-Lamp2 antibody (upper panel) and GAPDH protein level (lower panel) in mock transfected or pcDNA Lamp2c transfected Vero E6 Cells. D. Intracellular level of SRS CoV2 RNA (normalized to that of RP II) in Vero E6 cells transfected with the indicated plasmids and infected with SARS-CoV-2 for 48 hours. Data are mean ± SEM. E. Level of SARS-CoV-2 RNA (normalized to that of RP) in the culture medium of Vero E6 cells transfected with the indicated plasmids and infected with SARS-CoV-2 for 48 hours. Data are mean ± SEM. F. Western blot detection of Lamp1 protein level using Lamp1 antibody (upper panel) and GAPDH protein level (lower panel) in Huh7 cells transfected for 72 hours with non targeting SiRNA or Lamp1 SiRNA. G. Level of SARS-CoV-2 RNA (normalized to that of RP) in the culture medium of Huh7 cells transfected with Lamp1 SiRNA (72 hours) and infected with SARS-CoV-2 for 48 hours. Data are mean ± SEM. H. Level of SARS-CoV-2 RNA (normalized to that of RP II) in the culture medium of Huh7 cells transfected with Lamp1 SiRNA (72 hours) and infected with SARS-CoV-2 for 48 hours. Data are mean ± SEM.

In order to test if the observed reduction in viral RNA levels were specific to Lamp2, effect of Lamp1 depletion on SARS-CoV-2 replication was evaluated. Western blot using anti-Lamp1 antibody showed ∼90% reduction in Lamp1 protein level in siRNA transfected Huh7 cells at 48 hour and 72 hour time points (Fig 4F). QRT-PCR analysis of Lamp1 SiRNA treated SARS-CoV-2 infected cells did not show any significant change in the level of viral RNA in culture medium (Fig 4G) and intracellular samples (Fig 4H).

### Increased Lamp2 protein level in SARS-CoV-2 infected cells does not promote autophagolysosome formation

Lamp2a is the receptor for chaperone mediated autophagy and silencing of Lamp2 expression promoted SARS-CoV-2 replication. Therefore, we next measured the status of autophagy in SARS-CoV-2 infected cells. An increase in the level of LAMP2, LC3 II and p62 protein was observed in the SARS-CoV-2 infected Vero E6 and Huh7 cells (Fig 5A). No change in the level of Lamp1 was observed (Fig 5A). Treatment of SARS-CoV-2 infected Vero E6 and Huh7 cells with 3 methyl adenine (3-MA), which is a specific inhibitor of autophagy, completely diminished viral RNA in both culture medium and intracellular samples (Fig 5B-E). Next, Immunofluorescence staining of Lamp2 (green) and LC3 (Red) proteins was performed in SARS-CoV-2 infected or uninfected Vero E6 cells. Formation of autophagolysosome is indicated by colocalization of Lamp2 and LC3, which forms “yellow” color puncta in dual stained cells (35). No increase in autophagolysosome formation was visible in SARS-CoV-2 infected Vero E6 cells (Fig 5F).

**Figure 5.**
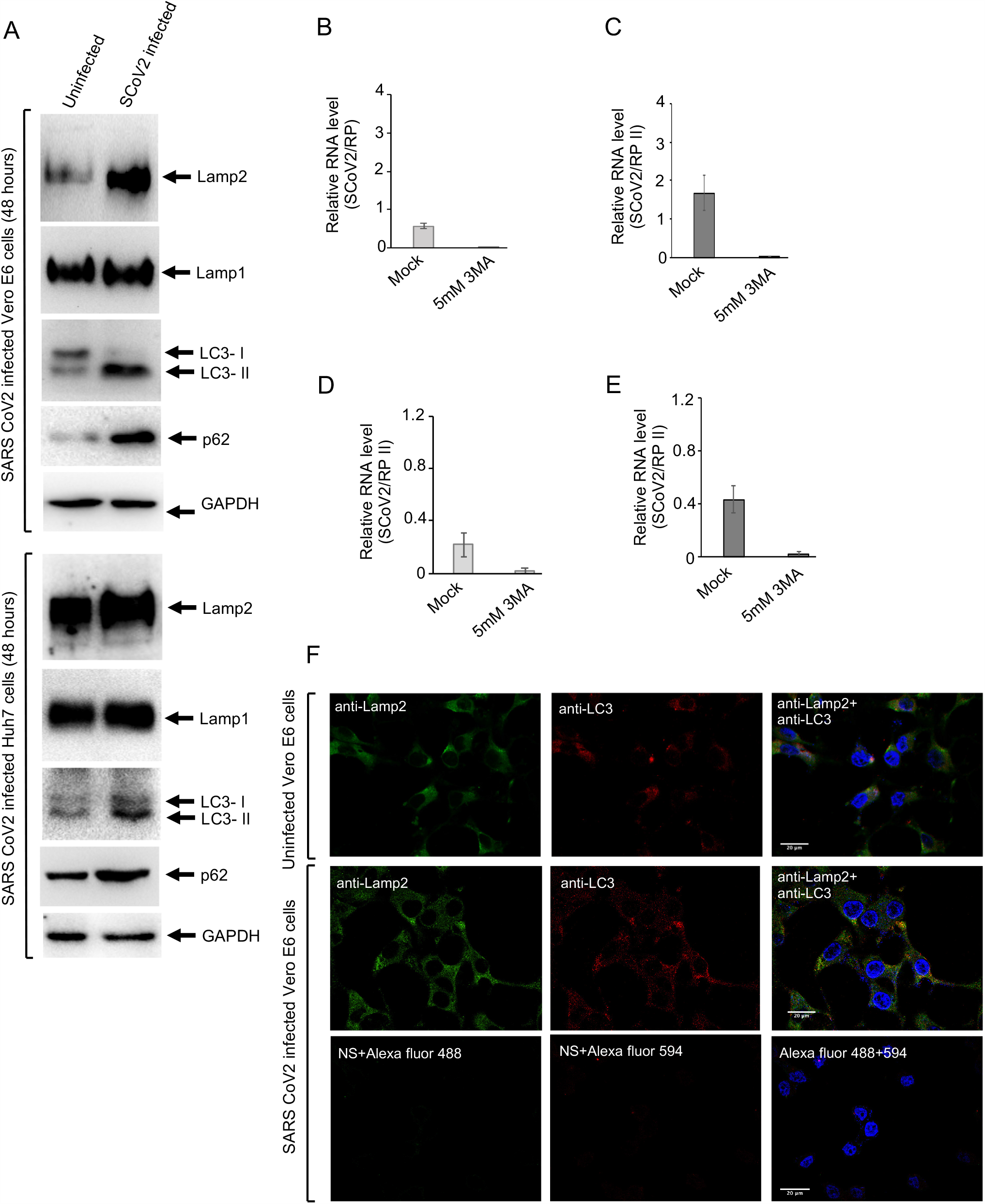
Increased Lamp2 and LC3-II levels in SARS-CoV-2 infected Vero E6 and Huh7 cells do not promote autophagolysosome formation. A. Western blot detection of indicated proteins in SARS-CoV-2 infected Vero E6 and Huh7 cells, 48 hour post-infection. B. Level of SARS-CoV-2 RNA (normalized to that of RP) in the culture medium of Vero E6 cells infected with SARS-CoV-2 and treated with 5mM 3MA for 48 hour. Data are mean ± SEM. C. Intracellular level of SRS CoV2 RNA (normalized to that of RP II) in Vero E6 cells treated with 5mM 3MA for 48 hour. Data are mean ± SEM. D. Level of SARS-CoV-2 RNA (normalized to that of RP) in the culture medium of Huh7 cells infected with SARS-CoV-2 and treated with 5mM 3MA for 48 hour. Data are mean ± SEM. E. Intracellular level of SRS CoV2 RNA (normalized to that of RP II) in Huh7 cells treated with 5mM 3MA for 48 hour. Data are mean ± SEM. F. Immunofluorescence staining of Lamp2 (green), LC3 (red), and nucleus (blue), in uninfected or 48 hour SARS-CoV-2 infected Vero E6 cells. Yellow indicates colocalization of the red and green signals. Scale bar is 20μM.

## Discussion

In this study, we identified 57 host proteins that interact with the 5’- and 3’-ends of the SARS-CoV-2 genomic RNA, combined those dataset with the PPI dataset proposed to be involved in viral replication and constructed an RPPI network, which is likely to be assembled during the replication of SARS-CoV-2. Interaction of 5’- and 3’-regulatory elements with virus-encoded and host proteins mediate their function. Since 5’- and 3’-regulatory elements are known to play indispensable roles in replication process of RNA viruses, identification of the components of the above-mentioned RNA-protein complex paves the way for designing novel antiviral therapeutic strategies against the SARS-CoV-2. Among the viral proteins, N protein of MHV, TGEV and SARS-CoV binds to the 5’-UTR (8). NSP8 and Nsp9 of MHV binds to the 3’-UTR (36). Many host proteins are known to bind to the 5’- and 3’-UTR of viral genome and regulate virus replication (7). During the course of this study, Schmidt *et al*. identified a number of host proteins that bind to SARS-CoV-2 RNA in infected Huh7 cells using the RAP-MS technique (23). Comparison of their dataset to that of this study identified 11 common proteins. This study identified additional 46 host proteins that associate with 5’- and 3’-UTRs of the SARS-CoV-2 RNA. This may be attributed to the ability of RaPID technique to detect both stable and transient as well as direct and indirect (as a protein complex) RNA-protein interactions. Specificity of the data is evident from the fact that only one protein (HIST1H3) is common between SARS-CoV-2 and HEV RPPI datasets. Note that similar to SARS CoV-2, HEV is also a positive strand RNA Virus, which contains a capped 5’-end, followed by a 5’-UTR and poly A tail preceded by a 3’-UTR (37). Therefore, some aspects of replication process might be similar in both viruses.

In addition to lungs, SARS-CoV-2 has been reported to cause enteric infection. High level of ACE2 is expressed in the intestinal epithelial cells and COVID-19 patients suffer from gastrointestinal disorders (4). Therefore, we hypothesized that host proteins that are crucial for the survival of the virus should be expressed in both lung and intestinal epithelium cells. Comparison of expression of 5’-300 and 3’-203 RNA binding proteins in lung and intestinal epithelium cells showed that 55 out of 57 proteins are expressed in both tissues, in support of the above statement.

Among the 5’- and 3’-UTR interacting host proteins, known drugs exist against RPL13, PDHA1, HABP4, MTHFD1, TUBA1A and PLA2G4A. Interestingly, some of the drugs being used/tested in COVID-19 patients such as Thalidomide and Ciclosonide act by targeting two of the above-mentioned proteins. Nevertheless, further studies are required to conclude if those drugs act in COVID-19 patients by targeting the observed RNA-protein interaction or through any other mechanism.

Gene ontology and Reactome pathway analysis of the SARS-CoV-2 5’- and 3’-UTR RPPI datasets show an enrichment of proteins involved in different processes such as translation initiation, RNA metabolism, infectious disease, which is expected. Therefore, the data obtained by the RaPID assay is an useful resource for future studies related to the SARS CoV-2. Comparison of SARS-CoV-2 5’-UTR and 3’-UTR interacting proteins obtained in the RaPID assay with those shown in other positive strand RNA viruses reveal a similar profile of many functional protein categories. Proteins belonging to the ribosome complex such as RPL7a, RPLP2, RPL13, RPL4 and translation initiation factor such as eIF5B were found to interact with the SARS-CoV-2 5’-and 3’-UTR. Being a capped RNA, SARS-CoV-2 genomic RNA is supposed to utilize the canonical cap-dependent translation process for producing viral proteins and above-identified host factors might be used by the virus for efficiently translating its own RNA. Heat shock proteins of the HSP70 and HSP90 family are required for maintenance of cellular homeostasis by virtue of their ability to recognize and fold or remove unfolded proteins (38). They are also important for proper assembly and/or disassembly of viral replication complex (39). They require the adapter protein STIP1 for optimal activity. In the case of SARS CoV-2, HSP71, HSP71L and STIP1 were found to interact with the 5’-UTR. Dead box RNA helicases such as DDX1 and DHX15 are known to associate with the TRS in MHV (7). DDX24 was identified to interact with the 3’-UTR of SARS CoV-2. DDX24 is known to associate with RNA and negatively regulate RIG-I-like receptor signaling, resulting in inhibition of host antiviral response (40). Several proteins encoded by the SARS-CoV-2 such as Nucleocapsid, ORF6 and ORF8 are known to inhibit host antiviral response (41). Interaction of DDX24 with SARS-CoV-2 3’-UTR might be an additional strategy to inhibit the host antiviral response. Further studies are required to confirm the above hypothesis.

ATP-binding cassette subfamily E member 1 (ABCE1) is another important host protein that interacts with the 5’-UTR. ABCE1 (RNAse L inhibitor) inhibits the activity of RNAse L, which is activated by the host antiviral response mechanism in response to RNA virus infection or IFN *α*/*β* stimulation (42). Active RNAse L cleaves the viral RNA, which is prevented in the presence of ABCE1. It will be interesting to explore the significance of 5’-UTR interaction with ABCE1. Note that ABCE1 was also identified as a direct interaction partner of SARS-CoV-2 RNA by Schmidt et al. (23). FKBP4 was also found to interact with the 5’-UTR. It is a member of the immunophilin protein family and binds to immunosuppressants FK506 and rapamycin (43). Further, Ras GTPase activating protein binding protein 2 (G3BP2), which is key to stress granules formation, was found to interact with the 5’-UTR of SARS CoV-2. G3BP1/2 has also been shown to bind to the N protein of SARS-CoV-2 (25). Stress granules play key roles controlling viral infections (44). N protein of SARS-CoV-2 is predicted to associate with the 5’-UTR. Hence, interaction of both N and 5’-UTR with G3BP2 may be an important strategy of SARS-CoV-2 to modulate the function of stress granules in infected cells. 5’-UTR was also found to interact with the exosome endoribonuclease RRP4, which is involved in cellular RNA processing and degradation. The RNA exosome complex is a quality control mechanism of the host that regulates mRNA turnover and degrades aberrant RNAs. SARS-CoV-2 might be modulating this pathway for its own benefit.

5’-UTR was also found to interact with Lamp2a, which is another key protein involved in quality control processes of the host. Lamp2a is the receptor for the chaperone mediated autophagy (45). Our data shows an increase in autophagic flux in the SARS-CoV-2 infected cells and significant reduction in viral RNA level by inhibition of autophagy. Therefore, it appears that increased autophagic flux promotes SARS-CoV-2 replication, as seen in many other RNA viruses (46). However, there was no change in autophagolysosome formation in SARS-CoV-2 infected cells. Similar phenomenon is observed in the case of HCV infection (47). Lack of Lamp2 protein increased the level of viral RNA in the SARS-CoV-2 infected cells. Therefore, Lamp2a interaction with the 5’-UTR is unlikely to play a role in promoting viral replication.

Lamp2b and Lamp2c isoforms are known to directly interact with RNA and DNA, leading to RNautophagy/DNautophagy (48). Lamp2c is predominantly involved in importing nucleic acids to lysosome for degradation (49). Lamp2 isoforms vary in their C-terminal region. Although SARS-CoV-2 5’-UTR interacts with Lamp2a, SiRNA used in our study acts by binding to the 5’ end of the Lamp2 transcript. Therefore, it acts on all Lamp2 variants. The antibody used in our study recognizes all Lamp2 isoforms. However, since Lamp2c overexpression did not affect viral RNA level, it is unlikely that the observed increase in viral RNA level in Lamp2 siRNA treated cells were attributed to a block in RNautophagy. Lamp2a-5’UTR interaction may be involved in delivering viral RNA to endosome, for recognition by TLR7, the sensor for ssRNA. Lamp2a and 5’-UTR interaction may also be involved in sequestering away Lamp2a from inducing chaperone mediated autophagy. At the same time, increased autophagic flux provides energy and prevents apoptosis of the infected cells, which provides a favorable environment to the virus to complete its life cycle. Further studies are required to understand the actual process. Nevertheless, the current study identifies the repertoire of host proteins that associate with the SARS-CoV-2 5’- and 3’-UTR and demonstrate a prosurvival role of the host Lamp2 protein against the Virus.

## Materials and Methods

### Plasmids and reagents

Sequence corresponding to 1-300 nucleotides at the 5’ end (5’-300) and 29676-29900 at the 3’end (3’-203) of the SARS CoV2 isolate Wuhan-hu-1 genome (Genbank: NC_045512) were commercially synthesized and cloned into pUC57 vector (Genscript, New Jersey, USA). RNA motif plasmid cloning backbone [(pRMB) (Addgene plasmid # 107253; http://n2t.net/addgene:107253; RRID:Addgene_107253)], BASU RaPID plasmid (Addgene plasmid # 107250; http://n2t.net/addgene:107250; RRID:Addgene_107250), RNA motif plasmid EDEN15 (Addgene plasmid # 107252; http://n2t.net/addgene:107252; RRID:Addgene_107252) were gifted by Paul Khavari. pCDNA Lamp2b was a gift from Joshua Leonard (Addgene plasmid # 71292; http://n2t.net/addgene:71292; RRID:Addgene_71292) and pCDNA lamp2c was a gift from Janice Blum (Addgene plasmid # 89342; http://n2t.net/addgene:89342; RRID:Addgene_89342). pcDNA Lamp2a (Cat no. HG29846-CY) was purchased from Sino Biologicals (Beijing, China). Anti-Lamp2 (Cat. No. 49067), anti-Lamp1 (Cat no. 9091), anti-LC3B (Cat no. 83506) and anti-P62 (Cat no. 8025s) antibodies were from Cell signalling Technology (Massachusetts, USA). Anti-GAPDH antibody (Cat No. SC-25778) was from Santacruz Biotechnology (Texas, USA). Anti-Flag antibody (Cat no. A190-101) was from Bethyl laboratory (Texas, USA). Anti-CUGBP1 antibody (Cat no. STJ92521) was from St John’s Laboratory, (London, UK). Goat anti-rabbit IgG-HRP (Cat No. 4030-05) and Goat anti-mouse IgG-HRP (Cat No. 1030-05) was from Southern Biotech (Alabama, USA). Goat anti-rabbit IgG Alexa fluor 488 (Cat no. A-11008), goat anti-mouse IgG Alexa fluor 647 (A-21235) and Antifade gold with DAPI (P36931) was from Thermo Fisher Scientific (Massachusetts, USA). 3-MA (Cat no. M9281) and Anti-HA antibody (Cat no. A190-101) was from Sigma (Missouri, USA). Nontargeting siRNA (Cat no. D-001810-10-20), Lamp1 siRNA (Cat no. L-013481-02-0005) and Lamp2 siRNA (Cat no. L-011715-00-0005) were from Dharmacon (Colorado, USA).

### Mammalian cell culture and transfection

Vero E6 and HEK 293T cells were obtained from ATCC (Virginia, USA). Huh7 cells have been described (50). Cells were maintained in Dulbecco’s modified Eagle medium (DMEM) containing 10% Fetal bovine Serum (FBS), 50 I.U./mL Penicillin and Streptomycin, in 5% CO_2_. For plasmid transfection, cells were seeded at 70-80% confluency in DMEM+10% FBS and incubated overnight. Next day, cells were transfected with desired plasmid DNA using Lipofectamine 2000 transfection reagent (Life Technologies, California, USA) at 1:1 ratio, following manufacturer’s instruction. 6-8 hours post-transfection, culture medium was replaced with fresh DMEM+10% FBS.

For experiments involving siRNA mediated gene silencing, Huh7 or Vero E6 cells were seeded at 70-80% confluency on 12 well TC dishes and incubated overnight at 37°C, 5% CO_2_. Next day, 25nmol siRNA was transfected into each well using 0.35 μl Dharmafect transfection reagent, following manufacturer’s instruction (Dharmacon, Colorado, USA). 18 hours post-transfection, culture medium was replaced with DMEM+10% FBS and cells were maintained at 37°C, 5% CO_2_ until further manipulation.

### RaPID assay

5’-300 and 3’-228 regions were PCR (Polymerase chain reaction) amplified from the pUC57 vector and cloned into RNA Motif plasmid cloning backbone vector (pRMB) at the BsmBI restriction site using the following primers. 5’-300 FP: ATTAAAGGTTTATACCTTCCCAGG, 5’-300 RP: GTTTTCTCGTTGAAACCAGGG, 3’-203 FP: AATCTTTAATCAGTGTGTAACA, 3’-203 RP: TTTTTTTTGTCATTCTCCTAAG. Positive clones were checked by restriction mapping and confirmed by DNA sequencing.

Production of 5’-300 and 3’-203 RNA from the pRMB constructs was verified by transfection of pRMB 5’-300 and pRMB 3’-203 plasmids along with pRMB vector into HEK293T cells followed by RT-PCR detection of the 5’-300 and 3’-203 RNA using above-mentioned primer pairs. Expression of Bir*A ligase in HEK293T cells upon transfection of BASU plasmid was verified by western blot (see Fig 1D). Time duration for biotin treatment were optimized by co-transfection of RNA motif plasmid EDEN15 (pRMB EDEN15) and BASU plasmid in 6:1 ratio using Lipofectamine 2000 (1:1 ratio) into HEK293T cell. 42hour post transfection, 200μM biotin was added to the media and cells were incubated for 6, 12 and 18 hours. At each time point, whole cell extract was prepared in 2X laemmli buffer and equal amount of protein was resolved by 10% SDS-PAGE, followed by western blot using anti-Biotin antibody (Fig 1E).

For verifying the interaction between EDEN 15 and CUGBP1 by RaPID assay, HEK293T cells were transfected with (pRMB EDEN15) and BASU plasmids (6:1 ratio) using Lipofectamine 2000 (1:1 ratio, Life Technologies, California, USA). 40 hours post-transfection, culture medium was replaced and 200μM Biotin was added. 18 hours later, cells were washed three times in PBS and lysed in prechilled RIPA buffer (150 mM NaCl, 1.0% NP-40, 0.5% sodium deoxycholate, 0.1% SDS, 50mM Tris, pH 8.0) supplemented with protease and phosphatase inhibitor cocktail. The lysate was centrifuged at 14,000 × g for 45 min at 4°C. Free biotin was removed by Macrosep Advance Spin Filter 3K MWCO 20 mL (Cat no. 89131-974, VWR, USA). Aliquots of the samples were stored at −80°C for use as input in western blot. Protein concentration in each sample was determined using BCA protein assay kit (Cat no. Thermo Fisher Scientific, Massachusetts, USA) and equal amount of proteins were incubated with “MyOne Streptavidin C1” magnetic beads (Cat no. 65002, Life Technologies, California, USA) on a rotator at 4°C for 16 hours. Samples were washed four times with washing buffer, 2X Laemmli buffer was added to the beads and incubated at 95°C for 45 min. Both input and pull down samples were resolved by SDS-PAGE, followed by western blot using anti-CUGBP1 and GAPDH antibodies. Goat anti-rabbit IgG HRP conjugated secondary antibody was used to detect the protein bands by chemiluminescence (Clarity ECL substrate, Cat no.170-5061, Biorad, California, USA).

For mass-spectrometry, clarified cell lysate was precipitated in acetone at −20°C for 10 min, followed by storage at −80°C for 20 mins. The precipitates were solubilized in 8M urea. The protein concentration was estimated by BCA protein assay kit (Thermo Fisher Scientific, Massachusetts, USA).

#### In-Solution Digestion and Peptide Separation

Equal amount of protein (10 mg) from each sample was treated with 10 mM DTT (56°C, 30 min) and alkylated with 20 mM Iodoacetamide (IAA) at room temperature, 60 min, dark). Trypsin (Cat no. T1426, Thermo Fisher Scientific, Massachusetts, USA) was added to the samples at 1:20 (w/w) ratio and incubated at pH 8, 37°C for 24 h. Next, 1% formic acid was added to the samples and peptides were desalted using a SecPak C18 cartridge (Cat no. WAT020515, Waters, Massachusetts, USA) and subsequently lyophilised in a Speed Vac. High capacity streptavidin agarose resins (Cat no. 20361, Thermo Fisher Scientific, Massachusetts, USA) were used to pull down the biotinylated peptides. The beads were washed in binding buffer (50mM Na_2_HPO_4_, 150mM NaCl; pH 7.2) before use.

Lyophilised peptides were solubilized in 1 mL PBS and incubated with 150 μL washed streptavidin agarose beads for 2 hour at room temperature. Beads were washed once in 1mL PBS, once in 1 mL washing buffer (5% acetonitrile in PBS) and finally washed once in ultrapure water. Excess liquid was completely removed from the beads, and biotinylated peptides were eluted by adding 0.3 mL of a solution containing 0.1% formic acid and 80% acetonitrile in water by boiling at 95°C for 5 min. A total of 10 elutions were collected and dried together in a Speed Vac. Enriched peptides were desalted with C18 tips (Thermo Fisher Scientific, Massachusetts, USA), and reconstituted with solvent A (2% (v/v) acetonitrile, 0.1% (v/v) formic acid in water) for LC-MS/MS analysis.

#### LC-MS/MS acquisition

LC−MS/MS experiments were performed using Sciex 5600^+^ Triple-TOF mass spectrometer coupled with ChromXP reversed-phase 3 μm C18-CL trap column (350μm × 0.5 mm, 120 Å, Eksigent, AB Sciex, Massachusetts, USA) and nanoViper C18 separation column (75 μm × 250mm, 3 μm, 100 Å; Acclaim Pep Map, Thermo Fisher Scientific, Massachusetts, USA) in Eksigent nanoLC (Ultra 2D plus) system. The binary mobile solvent system was used as follows: solvent C (2% (v/v) acetonitrile, 0.1% (v/v) formic acid in water) and solvent B (98% (v/v) acetonitrile, 0.1% (v/v) formic acid). The peptides were separated at a flow rate of 200 nl/min in a 60 min gradient with total run time of 90 min. The MS data of each condition was acquired in IDA (information-dependent acquisition) with high sensitivity mode. Each cycle consisted of 250 and 100 ms acquisition time for MS1 (m/z 350−1250 Da) and MS/MS (100– 1800 m/z) scans, respectively, with a total cycle time of ∼2.3s. Each condition was run in triplicate.

#### Protein Identification and Quantification

All raw files (.wiff) were searched using “ProteinPilot” software (version 4.5, SCIEX) with the Mascot algorithm, for protein identification and semi quantitation against the SwissProt_57.15 database (20266 sequences after Homo sapiens taxonomy filter). The search parameters for identification of biotinylated peptides were as follows: (a) trypsin as a proteolytic enzyme (with up to two missed cleavages); (b) peptide mass error tolerance of 20 ppm; (c) fragment mass error tolerance of 0.20 Da; and (d) carbamido-methylation of cysteine (+57.02146 Da), oxidation of methionine (+15.99492 Da), deamination of NQ (+0.98416) and biotinylation of lysine (+226.07759 Da) as variable modifications. Quality of data between different samples and replicates were monitored by Pearson-correlation plot of peptide intensity against each run.

#### Data analysis

Proteins with at least one corrected biotinylated peptide and ≥ 15 PEP score were considered to be identified successfully, extracted from the gaussian smooth curve. Next, all data were analyzed using a web based tool, Bioinformatics & Evolutionary Genomics (http://bioinformatics.psb.ugent.be/webtools/Venn/) to generate the venn diagram to identify proteins that are unique interaction partner of 5’-300 and 3’-203 RNA. The background subtracted dataset was sorted on the basis of following parameters (minimum 2 unique peptides and “Prot score” of 40 or more) to generate the final list of proteins, which were considered for further studies (see Table 1). The mass spectrometry proteomics data have been deposited to the ProteomeXchange Consortium via the PRIDE partner repository.

### Bioinformatics Analysis

RNA secondary structure was analysed using the “mfold” program (http://bioinfo.rpi.edu/applications/mfold/old/RNA), based on minimum free energy calculation at 25°C (51). The virus-host RPPI dataset was visualized using Cytoscape (version 3.1.0) (24). NetworkAnalyzer plugin in Cytoscape was used to compute the topological parameters and centrality measures of the network. Gene ontology “GO” and Reactome pathway analysis was performed using the “Gene Set Enrichment Analysis” tool [https://www.gsea-msigdb.org/gsea/index.jsp (26, 27)].

### SARS-CoV-2 infection

SARS-CoV-2 was obtained from BEI Resources (NR-52281, SARS-related coronavirus 2 isolate ISA-WA1/2020), amplified in Vero E6 cells in the BSL3 facility of THSTI, India, titrated and stored frozen in aliquots. For SARS-CoV-2 infection studies, 200 µl of stock virus was diluted in serum free media to 2000 TCID50/ml and added to Vero E6/ Huh7 cell monolayer (seeded in 24 well plate) for 1 hour at 37°C, supplemented with 5% CO_2_. One hour post-incubation, the infection medium was removed, cells were washed twice with 500 µl of serum free media and fresh DMEM+10% FBS was added. In case of plasmid over expression or siRNA transfection study, cells were transfected with respective DNA or siRNA 24 hours prior to infection with SARS-CoV-2. In case of 3-MA treatment, 5mM (final concentration) 3-MA was added to the culture medium during infection, again added to the complete medium after removing the infection medium and maintained for 48 hours, followed by collection of culture medium and cells and subsequent experiments.

### Fluorescence *In Situ* Hybridization (FISH)

#### Preparation of RNA probe

Sequence corresponding to the 5’-300 bases of SARS-CoV-2 was PCR amplified from pUC57 vector and cloned into the pJet1.2 vector (Thermo Fisher Scientific, Massachusetts, USA) under the control of T7 promoter. pJet1.2_5’-300 plasmid was linearized by restriction digestion with XbaI. DIG-11-UTP (Cat no. 11209256910, Roche, Basel) labeled RNA was *in vitro* synthesized using Maxi script *in vitro* transcription kit (Thermo Fisher Scientific, Massachusetts, USA), following manufacturer’s instruction. Template DNA was removed by treatment with Dnase I, followed by precipitation of RNA using LiCl. An aliquot of the probe was resolved by agarose gel electrophoresis to monitor its size and integrity.

#### Fluorescence in situ hybridization

FISH was done as described with minor modifications in which, Tyramide super boost kit (Cat No. B40933, Thermo Fisher Scientific, Massachusetts, USA) was used to detect the fluorescence signal (52). In summary, Vero E6 Cells were seeded at 70-80% confluency on coverslip overnight. Next day, cells were infected with the SARS-CoV-2 (see SARS-CoV-2 infection section) and incubated for 24 or 48 hours. Next, cells were washed three times in PBS, followed by fixation in 4% Paraformaldehyde for 10 min at room temp. Cells were rehydrated in 2xSSC buffer (300mM NaCl, 30mM Sodium Citrate) for 10 min at room temp, followed by incubation with 50% formamide in 2xSSC buffer for 30 min at room temp. Next, coverslips were washed four times with warm hybridization buffer (10% formamide in 2xSSC, warmed to 70°C), followed by incubation with hybridization buffer containing 2 ng/μl of probe for 3 days at 42°C in a humified incubator. Cells were washed two times with 4xSSC buffer for 10min each, at 42°C and incubated with 20ug/ml RNase A in 2xSSC buffer for 30min at 37°C. Next, cells were washed once with 2xSSC buffer and once with 0.1xSSC buffer for 10 min at 42°C. Endogenous peroxidase activity was quenched by adding 50 μl 3% H_2_O_2_ solution (provided in the Tyramide super boost kit) and incubating at room temp for 1 hour. Next, cells were blocked in 4% BSA (in PBS) for 1 hour at room temp, followed by incubation with anti-LAMP2 (1:50 dilution) and biotin conjugated anti-DIG antibody (1:200 dilution) prepared in 4% BSA in PBST (PBS+ 0.2% Tween-20) for overnight at 4°C. Next day cells were washed three times in PBS (10 min each) and incubated with HRP-conjugated streptavidin (provided in the Tyramide super boost kit) for 60 min at room temp. Next, cells were washed three times in PBS and incubated with Goat-anti rabbit IgG Alexa Fluor 488 (1:1000 dilution, prepared in 4% BSA+PBS+ 0.1% Tween 20) for 1 hour at room temp. Cells were washed three times in PBS. Next, freshly prepared Tyramide working solution was added to the cells and incubated for 5 min at room temp, followed by addition of stop solution (provided with the Tyramide super boost kit) and incubation for 3 min. Next, cells were washed three times in PBS and coverslips were mounted on slide using antifade gold. Images were acquired using a 100X objective in a confocal microscope (Olympus FV3000) and analysed by Image lab FIJI software.

### RNA isolation and quantitative real time PCR (QRT-PCR) assay

Intracellular RNA was isolated using TRI reagent (MRC, Massachusetts, USA), followed by reverse transcription (RT) using Firescript cDNA synthesis kit (Solis Biodyne, Estonia). RNA from culture medium was isolated using Qiagen viral RNA mini kit (Qiagen, Germany), followed by reverse transcription (RT) using Firescript cDNA synthesis kit (Solis Biodyne, Estonia). Random hexamers were used in cDNA synthesis. SYBR green based QRT-PCR was done as described earlier (50). Following primers were used: SCoV2 QPCR FP: 5’-TGGACCCCAAAATCAGCGAA, SCoV2 QPCR RP: 5’-TCGTCTGGTAGCTCTTCGGT; RP II FP: 5’-GCACCACGTCCAATGACAT, RP II RP: 5’-GTCGGCTGCTTCCATAA, RP FP: 5’-AGATTTGGACCTGCGAGCG, RP RP: 5’-GAGCGGCTGTCTCCACAAGT. Taqman based QRT-PCR was done as described earlier, following the protocol suggested by Centre for Disease Control, USA (53). Following primers and probes were used. N1 FP: 5’-GACCCCAAAATCAGCGAAAT, N1 RP: 5’-TCTGGTTACTGCCAGTTGAATCTG, N1 Probe: 5’-FAM-ACCCCGCATTACGTTTGGTGGACC-BHQ1, N2 FP: 5’-TTACAAACATTGGCCGCAAA, N2 RP: 5’-GCGCGACATTCCGAAGAA, N2 Probe: 5’-FAM-ACAATTTGCCCCCAGCGCTTCAG-BHQ1; RP FP: 5’-AGATTTGGACCTGCGAGCG, RP RP: 5’-GAGCGGCTGTCTCCACAAGT, RP Probe: 5’-FAM-TTCTGACCTGAAGGCTCTGCGCG-BHQ1. Absolute quantification was used in SYBR green QRT-PCRs. Standard plot was generated by serial dilution of known quantity of template. SCoV2 PCR values were normalized to that of RNA Polymerase II (RP II, intracellular RNA) or RNAse P (RP, culture medium RNA).

### Statistics

Data are represented as mean±SEM of three experiments. P values were calculated using two-tailed student t-Test (paired two sample for means).

### Immunofluorescence assay (IFA)

Confocal imaging was performed as described (50). In brief, Huh7 and Vero E6 cells were fixed with 4% Paraformaldehyde for 10 min at room temp, followed by incubation in blocking buffer (4% BSA in PBS) for 1 hour at room temp and incubation with Lamp2 and LC3b antibodies (1:50 and 1:50 dilution, respectively) in antibody dilution buffer (3% BSA in PBS+0.1% Tween-20) for 16 hours, at 4°C. Coverslips were washed three times in PBS, followed by incubation with 1:500 dilution of goat anti-rabbit Alexa Fluor 488 and 1:500 dilution of goat anti-mouse Alexa Fluor 647 antibodies (Thermo Fisher Scientific, Massachusetts, USA) in antibody dilution buffer at room temp for 1 hour. Coverslips were washed three times in PBS and mounted on glass slides using antifade gold reagent (Thermo Fisher Scientific, Massachusetts, USA). Images were acquired using a 60X objective in a confocal microscope (Olympus FV3000) and analysed by Image lab FIJI software.

### Western blot assay

Samples were resolved by SDS-PAGE, transferred to 0.4μ PVDF membrane. Membranes were blocked for 1 hour at room temp using 5% skimmed milk (in PBS). Next, membranes were incubated with primary antibody overnight in PBST (PBS+0.05% Tween-20) + 5% skimmed milk at 4°C. Blots were washed 3 times in PBST, followed by incubation with HRP-tagged secondary antibody at room temp for 1 hour. Blots were washed 3 times in PBST and protein bands were visualized by enhanced chemiluminescence using a commercially available kit (Bio Rad, California, USA).

## Acknowledgments

RNA motif plasmid cloning backbone (pRMB), BASU RaPID plasmid, RNA motif plasmid EDEN15 were kindly gifted by Dr Paul Khavari. pCDNA Lamp2b was a gift from Joshua Leonard and pCDNA Lamp2c was a gift from Janice Blum. SARS-Related Coronavirus 2, isolate USA-WA1/2020 (NR-52281) was deposited by the Centers for Disease Control and Prevention and obtained through BEI Resources, NIAID, NIH, USA. We thank Dr Tripti Shrivastava for generously providing the anti-N antibody (GeneTex Inc,USA). This study was funded by the Science and Engineering Research Board (SERB), Govt of India, IRHPA grant (IPA/2020/000233) to MS and THSTI core grant to MS. RV and SS are supported by a senior research fellowship from the Council of Scientific and Industrial Research, Govt of India, SK is supported by a senior research fellowship from the Department of Biotechnology, Govt of India.

## Legend to supplementary figures

**S1 Figure. Schematic of predicted secondary structure of SARS-CoV-2 5’-300 RNA**.

A. Schematic of predicted secondary structure of SARS-CoV-2 5’-300 RNA. “*” denotes an unannotated stem loop in the 5’-UTR.

B. Schematic of predicted secondary structure of SARS-CoV-2 5’-300 RNA + SL-A+ SL-B hybrid RNA.

**S2 Figure. Schematic of predicted secondary structure of SARS-CoV-2 3’-203 RNA**.

A. Schematic of predicted secondary structure of SARS-CoV-2 3’-203 RNA.

B. Schematic of predicted secondary structure of SARS-CoV-2 3’-203 RNA + SL-A + SL-B hybrid RNA.

**S3 Figure. Normalized distribution curve of LC-MS-MS identified peptides**.

A. PEP score wise normal distribution of all peptides identified in LC-MS-MS in different biological replicates.

**S4 Figure. RPPI network of SARS-CoV-2 5’-300 RNA + 3’-203 RNA + PPI involved in virus replication**.

A. Schematic of RPPI network of the 5’-300+3’-203 SARS-CoV-2 RNAs + PPI involved in virus replication. Black nodes: 5’-300 or 3’-203; Blue or red nodes: Host protein. Host proteins that interact with each other contain yellow or green color inside the node. Common host proteins that interact with both 5’-3000 and 3’-203 RNA are represented with dual (red+blue) color. Viral protein nodes are labeled. Host proteins involved in PPI with viral proteins are indicated in sky blue. Interaction among viral proteins is represented by blue lines.

B. Schematic of PPI network involved in SARS-CoV-2 replication. Viral protein nodes are labeled. Host proteins involved in PPI with viral proteins are indicated in sky blue. Interaction among viral proteins is represented by blue lines.

**S5 Figure. GO analysis of 5’-300 and 3’-203 SARS-CoV-2 RNA interacting proteins**.

Proteins involved in top 10 biological processes are indicated in blue.

**S6 Figure. Reactome pathway analysis of 5’-300 and 3’-203 SARS-CoV-2 RNA interacting proteins**.

Proteins involved in top 10 pathways are indicated in blue.

**S7 Figure. Lamp2 modulates viral RNA level in SARS-CoV-2 infected Huh7 cells**.

A. Level of SARS-CoV-2 RNA [normalized to that of RNase P (RP II)] in the culture medium (secreted) and inside Huh7 cells (intracellular) that are infected with the SARS-CoV-2 for the indicated periods. Real-time PCR reactions were performed using SYBR green based protocol. Data are mean ± SEM.

B. Western blot detection of SARS-CoV-2 nucleocapsid protein (upper panel) and GAPDH (lower panel) in SARS-CoV-2 infected Huh7 cells, 48 hour post-infection.

C. Western blot detection of Lamp2 (Lamp2 antibody, upper panel) and GAPDH lower panel) protein in Huh7 cells transfected for 72 hours with NT SiRNA or Lamp2 siRNA.

D. Level of SARS-CoV-2 RNA (normalized to that of RP) in the culture medium of Huh7 cells transfected with Lamp2 SiRNA and infected with SARS-CoV-2 for 48 hours. Data are mean ± SEM.

E. Intracellular level of SARS-CoV-2 RNA (normalized to that of RP II) in Huh7 cells transfected with Lamp2 siRNA and infected with SARS-CoV-2 for 48 hours. Data are mean ± SEM.

F. Western blot detection of Lamp2a protein level [using anti-HA antibody (upper panel) and anti-Lamp2 antibody (middle panel)] and GAPDH protein level (lower panel) in mock transfected or pcDNA Lamp2a transfected Huh7 Cells.

G. Western blot detection of Lamp2b protein level [using anti-HA antibody (upper panel) and anti-Lamp2 antibody (middle panel)] and GAPDH protein level (lower panel) in mock transfected or pcDNA Lamp2b transfected Huh7 Cells.

H. Western blot detection of Lamp2c protein level using anti-Lamp2 antibody (upper panel) and GAPDH protein level (lower panel) in mock transfected or pcDNA Lamp2c transfected Huh7 Cells.

I. Level of SARS-CoV-2 RNA (normalized to that of RP) in the culture medium of Huh7 cells transfected with indicated plasmids and infected with SARS-CoV-2 for 48 hours. Data are mean ± SEM.

J. Intracellular level of SRS CoV2 RNA (normalized to that of RP II) in Huh7 cells transfected with indicated plasmids and infected with SARS-CoV-2 for 48 hours. Data are mean ± SEM.

## Supporting information

S1 Fig. Schematic of predicted secondary structure of SARS-CoV-2 5’-300 RNA.

S2 Fig. Schematic of predicted secondary structure of SARS-CoV-2 3’-203 RNA.

S3 Fig. Normalized distribution curve of LC-MS-MS identified peptides.

S4 Fig. RPPI network of SARS-CoV-2 5’-300 RNA + 3’-203 RNA + PPI involved in virus replication.

S5 Fig. GO analysis of 5’-300 and 3’-203 SARS-CoV-2 RNA interacting proteins.

S6 Fig. Reactome pathway analysis of 5’-300 and 3’-203 SARS-CoV-2 RNA interacting proteins.

S7 Fig. Lamp2 modulates viral RNA level in SARS-CoV-2 infected Huh7 cells.

S1 Table. List of host proteins that associate with SARS-CoV-2 5’-300 and 3’-203 RNA.

S2 Table. Host proteins that interact with the 3’-end of HEV genome, identified by RaPID assay.

S3 Table. Comparison of expression of prey proteins identified in RaPID assay in human Lungs and Intestine.

